# Open Science Datasets from PREVENT-AD, a Longitudinal Cohort of Pre-symptomatic Alzheimer’s Disease

**DOI:** 10.1101/2020.03.04.976670

**Authors:** Jennifer Tremblay-Mercier, Cécile Madjar, Samir Das, Alexa Pichet Binette, Stephanie O.M. Dyke, Pierre Étienne, Marie-Elyse Lafaille-Magnan, Jordana Remz, Pierre Bellec, D. Louis Collins, M. Natasha Rajah, Veronique Bohbot, Jeannie-Marie Leoutsakos, Yasser Iturria-Medina, Justin Kat, Richard D. Hoge, Serge Gauthier, Christine L. Tardif, M. Mallar Chakravarty, Jean-Baptiste Poline, Pedro Rosa-Neto, Alan C. Evans, Sylvia Villeneuve, Judes Poirier, John C. S. Breitner, the PREVENT-AD Research Group

## Abstract

To move Alzheimer Disease (AD) research forward it is essential to collect data from large cohorts, but also make such data available to the global research community. We describe the creation of an open science dataset from the PREVENT-AD (PResymptomatic EValuation of Experimental or Novel Treatments for AD) cohort, composed of cognitively unimpaired older individuals with a parental or multiple-sibling history of AD. From 2011 to 2017, 386 participants were enrolled (mean age 63 years old ± 5) for sustained investigation among whom 349 have retrospectively agreed to share their data openly. Repositories are findable through the unified interface of the Canadian Open Neuroscience Platform (https://portal.conp.ca/) and contain up to five years of longitudinal imaging data, cerebral fluid biochemistry, neurosensory capacities, cognitive, genetic, and medical information. Imaging data can be accessed openly at https://openpreventad.loris.ca while most of the other information, sensitive by nature, is accessible by qualified researchers at https://registeredpreventad.loris.ca. In addition to being a living resource for continued data acquisition, PREVENT-AD offers opportunities to facilitate understanding of AD pathogenesis.

## BACKGROUND AND SUMMARY

Dementia is the final stage of Alzheimer’s disease (AD), representing the culmination of a process that begins decades before onset of symptoms [1–3]. Characterizing and tracking the pre-symptomatic stage of AD requires methods sensitive to the disease’s early manifestations. These may include not only subtle cognitive decline, but also biochemical changes and structural or functional brain alterations. Studying these pre-symptomatic changes is crucial to a full understanding of AD, and their precise measurement is critical for trials of interventions that seek to prevent symptom onset.

To meet this challenge, in 2010 investigators at McGill University and the Douglas Mental Health University Institute Research Centre created a Centre for **St**udies **o**n **P**revention of **A**lzheimer’s **D**isease (StoP-AD Centre). The Centre’s prime objective was to pursue innovative studies of pre-symptomatic AD, with efforts to provide relatively enriched samples for prevention trials requiring individuals at-risk of developing the disease [4]. To this end, the StoP-AD Centre developed an observational cohort for **PR**e-symptomatic **EV**aluation of **E**xperimental or **N**ovel **T**reatments for **AD**(PREVENT-AD). To increase the probability that participants would harbor the earliest changes associated with pre-symptomatic AD, entry criteria required intact cognition and a parental or multiple-sibling family history of AD. It is well-documented that populations with such a family history of AD-like dementia have a 2-3 fold relative increase in risk of AD dementia [5, 6].

The cohort was genotyped and followed by naturalistic studies of cognition, neurosensory capacities, cerebrospinal fluid (CSF) biochemistry, magnetic resonance neuroimaging (MRI) and by medical and clinical evaluations. The goal was to test a vast array of well-known biomarkers of AD pathology (amyloid and tau level in CSF) and neurodegeneration (cortical thickness and volume), but also to include more experimental measurements (e.g. episodic memory task fMRI) and other promising biomarkers (e.g. neurosensory measures). Precise measurement of the biomarkers mentioned above was required not only to monitor the progression of asymptomatic AD, but also to assess the effects of preventive interventions years before symptom onset. The main StoP-AD Centre clinical trial investigated the impact of naproxen sodium, a non-steroid anti-inflammatory drug, on the trajectory of a composite score of AD biomarkers (INTREPAD trial - NCT0270281). Data collection protocols remained the same between 2011 and 2017 with only a few additions. The data repositories and related methods described here refer to the data collected at the StoP-AD Centre, in the observational cohort and the INTREPAD trial from November 2011 to November 2017, the so-called PREVENT-AD ‘Stage 1’. During this interval, a total of 425 participants completed their baseline visit (BL), with 195 participants initially enrolled in the INTREPAD trial. A total of 386 met criteria for sustained investigation and 349 retrospectively agreed to share their data openly (Figure 1).

**FIGURE 1:**
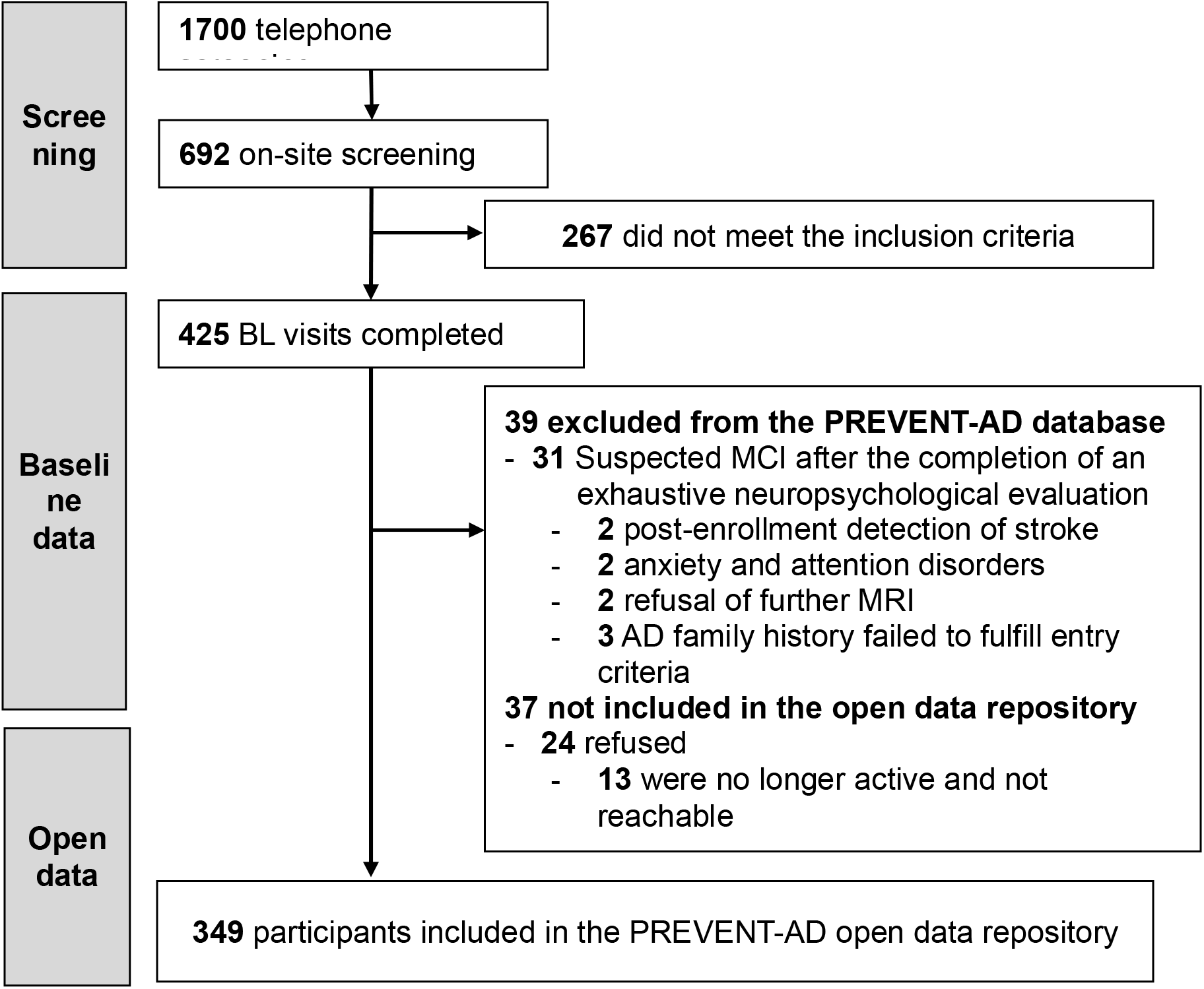
PREVENT-AD candidates enrolled between November 2011 and November 2017. 425 participants performed a baseline visit (BL) among which the datasets of 386 participants were shared with internal collaborators for analysis. 349 of these participants agreed to have their data openly shared, while one of them specifically refused to share data in the registered repository. MCI=Mild Cognitive Impairment, MRI = Magnetic Resonance Imaging

PREVENT-AD data are now broadly available as an open science resource [7–9], providing opportunities for a larger number of researchers to analyze this rich dataset. Because the original PREVENT-AD ethics approval and consent process did not fully address open science data sharing plans, several steps related to ethics were required and are described below. Storage, management, quality control (QC), validation and distribution of PREVENT-AD data were performed in LORIS, a system designed for linking heterogeneous data (e.g. behavioral, clinical, imaging, genomics) within a longitudinal context [10, 11]. Up to 5 years of longitudinal follow-up are shared as part of this Stage 1, which includes a total of 1300 cognitive evaluations, 476 CSF sample analyses, 1559 MRI sessions and 1283 neurosensory tests, from 349 consenting participants.

As this research program is evolving, new biomarkers are being added. In February 2018, the data collection regimen was notably modified after a short period of inactivity and is now focused mainly on longitudinal cognitive, behavioral and lifestyle assessments as well as new neuroimaging modalities. These additional variables are described in the PREVENT-AD ‘Stage 2’ companion paper. (Pichet Binette et al., in preparation).

The multi-modal approach, the depth of the neurocognitive phenotyping of this population at increased risk of AD dementia along with the longitudinal nature of this single-site study render this dataset exceptional. The methods section will describe the PREVENT-AD cohort characteristics, data acquisition methods, and the approach used to create the data repositories for dissemination to the wider research community. Additional information about the PREVENT-AD program can be found at http://prevent-alzheimer.net/.

## METHODS

### 1. Development of Open Science data sharing

Making the PREVENT-AD dataset available for open science was a multi-step process achieved over approximately 2 years. The steps required to prepare the dataset before its dissemination on an open science platform included: ethical considerations (discussed below), data preparation in a structured and standardized way, dataset documentation (data dictionary, label convention, etc) and development of two LORIS databases for 1) Open and 2) Registered Access data dissemination. Significant efforts were needed to obtain additional ethics approval for the open science plans and proceed with a re-consent process for all participants. Most of the participants had remained actively involved in StoP-AD Centre activities, and this greatly facilitated the re-consent process. We also attempted to contact participants who were no longer associated with the Centre. We failed to reach only 13 participants out of 386 with data potentially to share. Even though partially de-identified data had been prepared for sharing with collaborating research teams, additional dataset de-identification steps were required to share data with a much larger community of researchers. For example, all brain images were “scrubbed” to remove any potentially identifying fields from their headers, and structural imaging modalities were defaced to prevent facial re-identification using 3D rendering [12]. Details about the procedure are described in the Data Record section. The initial PREVENT-AD study codes were assigned a new “public” alphanumeric code (e.g.: CONP00000), to which a participant’s identity cannot directly be linked; the ability to do so remaining exclusively with the StoP-AD team.

Decisions were made by PREVENT-AD investigators regarding the choice of variables to be shared (based on quality, reliability, and level of standardization) and their level of access (Open or Registered Access, based on data sensitivity and the risk of potential re-identification or misuse). Datasets were prepared for two different LORIS platforms depending on the level of access. *Open data* are available to anyone who requests an account, whereas a broader dataset is available through *Registered Access,* available only to *bona fide* researchers [13]. Registration is approved by the StoP-AD team upon verification of the applicant’s account information. Both repositories are discoverable through the unified interface of the Canadian Open Neuroscience Platform (CONP). Although costly, we expect that these efforts to prepare shared resources will increase the rate of scientific discovery in dementia research; this being our ultimate hope for people living with AD and their families.

### 2. Eligibility and enrollment assessments

A study nurse conducted preliminary eligibility screening over the phone or via an online questionnaire. Participants had to be 60 years of age or older, with an exception that individuals between 55 - 59 years old were eligible if their own age was within 15 years of symptom onset of their youngest-affected first-degree relative. Participants’ family history of “AD-like dementia” was ascertained either by a compelling AD diagnosis from an experienced clinician or, if such a report was not available, by use of a structured questionnaire developed for the Cache County Study [5]. The questionnaire was intended to establish memory or concentration issues sufficiently severe to cause disability or loss of function, having an insidious onset and gradual progression (as opposed to typical consequences of a stroke or other sudden insult).

An on-site eligibility visit (visit label EL00) then included more specific questions on family history of AD dementia, medical and surgical history, pharmacological profile, lifestyle habits, as well as physical and neurological examinations, blood and urine sampling. The blood sample was used for genotyping (see section 3.1) only after an individual was declared eligible to the program. The CAIDE score (Cardiovascular Risk Factors, Aging, and Incidence of Dementia risk score) was derived using data collected at entry into the program (age, sex, education, systolic blood pressure, body mass index (BMI), cholesterol, physical activity and APOE ε4 status) [14]. Two cognitive screening instruments assessed integrity of cognition: the Montreal Cognitive Assessment (MoCA) and the Clinical Dementia Rating (CDR) Scale [15, 16] including its brief cognitive test battery. When cognitive status was in doubt (MoCA typically ≤ 26/30 or CDR >0), a complete evaluation (2.5 hours of testing) was performed by a certified neuropsychologist. The aim of this assessment was to determine if the cognitive deficits detected by the screening tests fell within the range of mild cognitive impairment (MCI), did not meet MCI criteria or were simply circumstantial, see section ‘Management of cognitive decline’ for more details.

Subsequently, during the enrollment visit (visit label EN00), a ~30-minute Magnetic Resonance Imaging (MRI) session was acquired to rule out structural brain disease, while simultaneously ensuring participants’ familiarity with the MRI environment. Handedness was determined using the Edinburgh Handedness Inventory [17], and an electrocardiogram was performed. Enrollment also required further documentation of stable general health, availability of a study partner to provide information on daily functioning, and willingness to comply with study protocols (Table 1 for detailed inclusion/exclusion criteria). Specific INTREPAD clinical trial inclusion/exclusion criteria are in the publication describing results of the trial [18]. In brief, they were similar except for additional criteria related to gastrointestinal tract problems and specific contraindicated concomitant medication. Final determination of eligibility for PREVENT-AD program was made by clinical consensus between one or more study physicians, a research nurse, and a neuropsychologist. All consent procedures fulfilled modern requirements for human subjects’ protection, while avoiding excess participant burden. Consent forms were carefully crafted to use simple but comprehensive language (typically at an 8th grade reading level). Protocols, consent forms and study procedures were approved by McGill Institutional Review Board and/or Douglas Mental Health University Institute Research Ethics Board. Specific consent forms were presented prior to each experimental procedure.

**Table 1:** Inclusion and Exclusion Criteria.

|  |
| --- |
| **Inclusion criteria** |
| → Self-reported parental or multiple-sibling (2 or more*) history of Alzheimer-like dementia<br>→ Age 60 years or older (persons aged 55-59 years and $\leq 15$ years younger than their affected index relative were also eligible)<br>→ Minimum of 6 years of formal education<br>→ Study partner available to provide information on cognitive status<br>→ Sufficient fluency in spoken and written French and/or English<br>→ Ability and intention to participate in regular visits<br>→ Agreement for periodic donation of blood and urine samples<br>→ Agreement to participate in periodic multimodal assessments via MRI and LP for CSF collection (LP optional at first, then mandatory (in 2017) for participation)<br>→ Agreement to limit use of medicines as required by clinical trial protocols, if applicable<br>→ Provision of informed consent of the different protocols |
| **Exclusion criteria** |
| → Cognitive disorders - Known or identified during eligibility assessments (MoCA and CDR or exhaustive neuropsychological evaluation when needed)<br>→ Use of acetyl-cholinesterase inhibitors including tacrine, donepezil, rivastigmine, galantamine<br>→ Use of memantine or other approved prescription cognitive enhancer<br>→ Use of vitamin E at greater than 600 i.u. / day or aspirin at $>325$ mg / day<br>→ Use of opiates (oxycodone, hydrocodone, tramadol, meperidine, hydromorphone)<br>→ Use of NSAIDs or regular use of systemic or inhalation corticosteroids<br>→ Clinically significant hypertension (accepted if controlled medically), anemia, significant liver or kidney disease<br>→ Concurrent use of warfarin, ticlopidine, clopidrogel, or similar anti-coagulant<br>→ Current plasma Creatinine $>1.5$ mg/dl (132 mmol/l)<br>→ Current alcohol, barbiturate or benzodiazepine abuse/dependence |

**Inclusion and exclusion criteria for the PREVENT-AD observational cohort.** INTREPAD trial inclusion/exclusion criteria are specified in the publication describing the results (18). \*8 participants had only 1 sibling affected with AD-like dementia. Refer to 'List\_participants\_family\_history\_1\_sibling\_v1.0.txt' file. MRI: magnetic resonance imaging; LP: lumbar puncture; CSF: cerebrospinal fluid; MoCA: Montreal Cognitive Assessment; CDR: Clinical Dementia Rating; NSAID: non-steroidal anti-inflammatory drug;

### 3. Data collection overview

Eligible participants were enrolled either in the observational cohort or in the INTREPAD trial as both started enrollment around the same time. For both groups, the first annual visit is called baseline, labeled BL00, and follow-up (FU) visits are labeled FU12, FU24, FU36 and FU48, corresponding to the number of months after the baseline visit. Telephone follow-ups were conducted between on-site annual visits to keep contact with the participant and to update clinical information. Participants in the INTREPAD trial had more frequent on-site visits and telephone follow-ups for safety purposes. See Figure 2 for the data collection timelines of the observational and trial cohorts.

**FIGURE 2:**
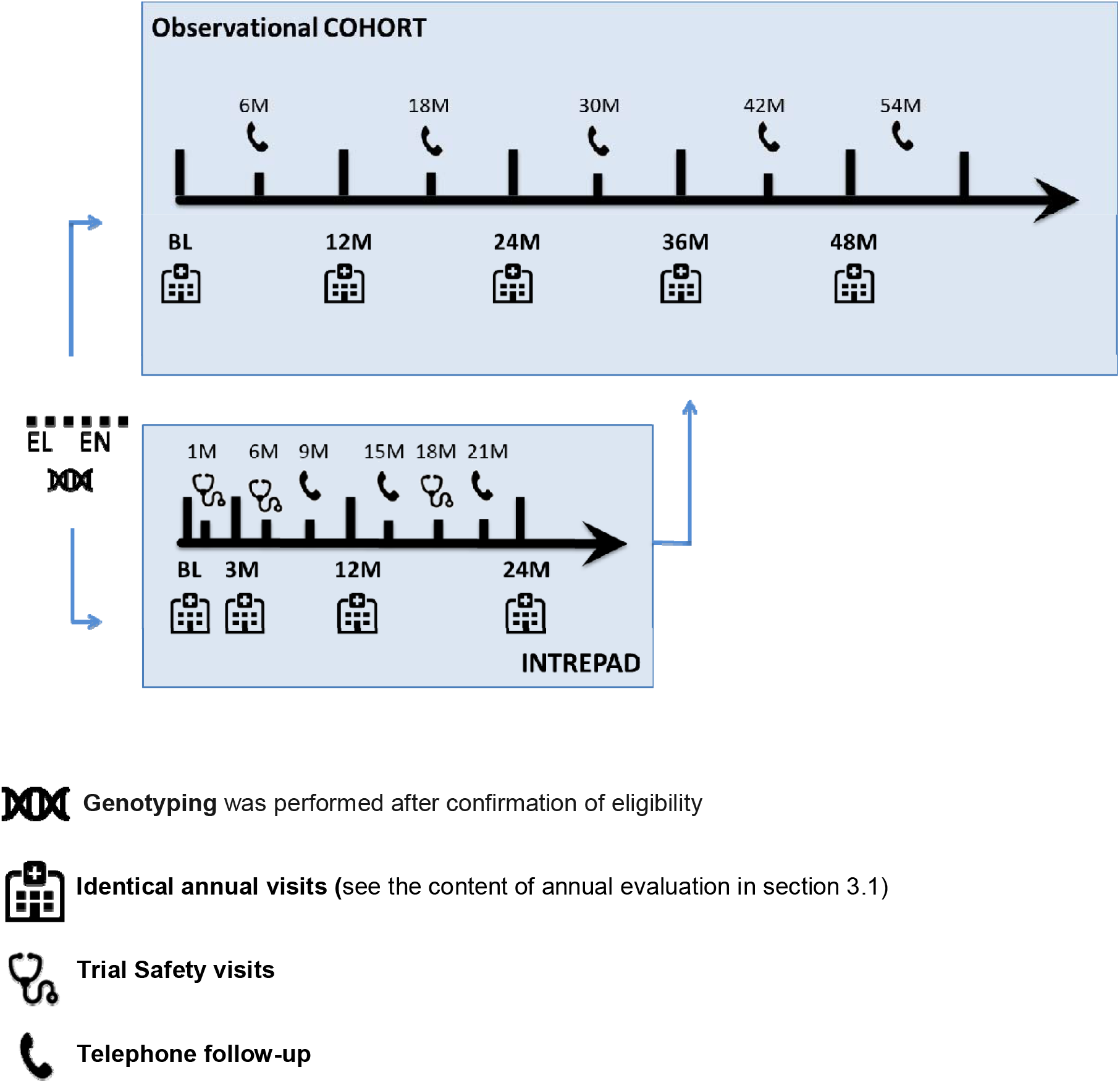
Timelines of observational cohort, INTREPAD trial. EL: eligibility visit; EN: enrolment visit; BL: Baseline visit; M: months;

#### 3.1 Annual evaluations

During each longitudinal visit (BL00, FU12, FU24, FU36 and FU48) for both observational and trial cohorts, a standardized cognitive evaluation, neurosensory tests, and an MRI scanning session (1 to 1.5 hours) were performed. On a separate day, participants who consented to the procedure donated CSF samples via lumbar puncture (LP). Medical conditions, pharmacological profile and various in-house health questionnaires were updated annually while blood and urine sampling, neurological and physical examinations were also performed. Routine laboratory results were obtained from a central medical laboratory in the Montreal area while ten milliliters of blood were centrifuged, aliquoted (plasma and red blood cells) and stored for further analysis in Dr. J. Poirier’s laboratory.

Details about each experimental procedure are described in section 4.

#### 3.2 INTREPAD trial

**INTREPAD** (***I**nvestigation of **N**aproxen **TR**eatment **E**ffects in **P**re-symptomatic **A**lzheimer’s **D**isease;* clinicaltrials.gov - NCT02702817) was a randomized, double-blinded, placebo-controlled two-year trial of low dose naproxen sodium (220 mg b.i.d.) conducted in 195 PREVENT-AD participants. Trial recruitment began in March 2012 and ended in March 2015. Treatment (active or placebo) duration was 24 months. Standard annual PREVENT-AD evaluations were supplemented with an additional identical session three months after randomization (FU03). The 3-month assessment was intended to determine whether treatment-related changes, if any, occurred gradually or had rapid onset. It also served as a run-in period, with a modified Intent-to-Treat analysis design that considered only those participants who remained on treatment through this initial interval. The primary outcome of the trial was a composite Alzheimer Progression Score (APS) derived using item response theory from various cognitive and biomarker measures [19]. For other trial data (such as study drug compliance, adverse events, etc), please refer to our results paper (Meyer *et. al.*, 2019) [18].

#### 3.3 Summary of the Open Science sub-sample

Among the 425 participants who underwent baseline visits, 386 were confirmed as appropriate for final data analysis (see Figure 1 for reasons for exclusion of the 39 others). From these, 349 participants (90.4 %) consented retrospectively to have their data included in the shared data repositories under the principles of open science (one participant specifically asked to only share data in the open repository leaving a number of 348 in the registered repository). Ethnicity and genetic background of the cohort are relatively homogeneous. The majority of participants in the shared dataset come from the greater Montreal area in Québec, Canada; 98.9% are Caucasian and 86% have French as mother tongue. Women are somewhat over-represented (102 men, 247 women) while the proportion of APOEL4 carriers (4/4 = 2.0%; 4/3 = 32.2%; 4/2 = 4.3%) is slightly higher than the general Caucasian population [20], in keeping with participants’ family history of AD. PREVENT-AD participants are on average, younger than those in most aging or AD studies, (mean age at baseline=63.6 ± 5.1 years old), are highly educated (15.4 ± 3.3 years of education) and cognitively unimpaired (MoCA score of 28.0 ± 1.6 out of 30). Up to four years of follow-ups are available (median 36-months of follow-up, IQR 36). Twelve-month data (FU12) are available for 278 participants, 24-month data (FU24) are available for 236 participants, 36-month (FU36) data are available for 177 participants and 48-month (FU48) data are available for 116 participants. Three-month (FU03) are available for 150 INTREPAD participants.

### 4. Data collection methods (related to shared data only)

#### 4.1 Genotyping

DNA was isolated from 200 μl whole blood using a QIASymphony apparatus and the DNA Blood Mini QIA Kit (Qiagen, Valencia, CA, USA). The standard QIASymphony isolation program was used following the manufacturer’s instructions. Allelic variants of seven genes associated with AD [21–26] (*APOE*: rs429358 and rs7412, *BDNF*: rs6265, *HMGCR*: rs3846662 *BCHE*: rs1803274, *TLR4*: rs4986790, *PPP2R1A*: rs10406151, *CDK5RAP2*: rs10984186) were determined using pyrosequencing (PyroMark24 or PyroMark96) or DNA microarray (Illumina) and are shared in the registered repository.

#### 4.2 Cognition

Cognitive performance over time was assessed using the ~30 minute **R**epeatable **B**attery for **A**ssessment of **N**europsychological **S**tatus (RBANS) [27] at baseline and each subsequent follow-up visit. This battery consists of 12 subtests (list learning, story learning, figure copy, line orientation, picture naming, semantic fluency, digit span, coding, list recall, list recognition, story recall, figure recall) that yield 5 Index scores (immediate memory, delayed memory, language, attention and visuospatial capacities) and a total score. The battery is available in both French and English in 4 equivalent versions to reduce practice effects in longitudinal assessment. The data were scored following the RBANS manual, which results in age-adjusted Index scores with a mean of 100 and standard deviation of 15. Subtest scores have a mean of 10 and standard deviation of 3. Additionally, we scored all participants using norms specified for individuals aged 60-69 years, thereby allowing detection of potential decline with advancing age. Both scores (age-adjusted and graded exclusively using age 60-69 norms) are available in the registered data repository.

At these annual visits, we also administered the AD8 questionnaire to the study partner (a family member or friend in regular contact with the study participant). The AD8 comprises 8 questions evaluating changes in the participant’s memory and functional abilities, and is intended to discriminate normal aging from very mild dementia [28]. AD8 total score and answers to each question are shared in the registered repository.

##### Management of cognitive decline

Once the MoCA and CDR performed at eligibility confirmed that the research participants were cognitively intact at entry, we performed the baseline RBANS and followed their cognition annually (+ 3-month FU in INTREPAD trial). However, at each visit, if the cognitive test results were lower than expected and the cognitive status was in doubt (MoCA<26 or CRD>0 (for screening tests at eligibility) or RBANS index score > 1 SD below the mean in two different cognitive domains (for the follow-ups)), a complete cognitive evaluation was requested by the study physician and was performed by a certified neuropsychologist. Suspicion of probable mild cognitive impairment (MCI) after this neuropsychological evaluation led to the exclusion (or ineligibility) of the research participant from the research program and referral to an affiliated memory clinic, as needed. This procedure was implemented to ensure our cohort was purely asymptomatic. An exception was made for participants from the INTREPAD trial since we needed to monitor potential adverse events, so INTREPAD participants showing cognitive decline were invited to pursue their annual visit at our Center. Notably, a significant portion of the comprehensive neuropsychological evaluations, triggered by low test results, turned out to be reassuring and did not reveal any cognitive deficits. In these cases, the participants were invited to continue their annual follow-up in our cohort as their low scores were considered ‘circumstantial’. From 2016, an extension to the PREVENT-AD protocol was approved to allow the follow-up of PREVENT-AD participants who developed MCI or dementia. Thus, the time point of conversion to probable MCI is documented in the data sharing repository and data related to this conversion point are also provided. This is described in more detail in the Stage 2 data sharing companion paper, in preparation.

#### 4.3 Neurosensory

##### 4.3.1 Smell identification

Odor identification (OI) abilities were tested in a 30-minute session in a well-ventilated room, using the standardized University of Pennsylvania Smell Identification Test (UPSIT) [29]. This test uses “scratch-and-sniff” stimuli of 40 items (4 randomized booklets of 10 odorants each). Although the test can be self-administered, a trained examiner administered the test to improve reliability. The UPSIT was administered at baseline and each follow-up visit, and total score and selected data are shared in the registered repository. Additional information on the use of the UPSIT in PREVENT-AD and related results are detailed in two publications [30, 31].

##### 4.3.2 Auditory processing

Central auditory processing (CAP) evaluations were added to the study in 2014. This instrument is therefore not available at all time points for every participant and was available only in French. We used both the Synthetic Sentence Identification with Ipsilateral Competing Message (SSI-ICM) test and the Dichotic Stimulus Identification (DSI) test [32, 33]. After having first been assessed for simple auditory acuity (with monosyllabic words), participants were asked to identify spoken “pseudo-sentences,” either with various sound levels of a distracting background narrative (SSI-ICM) or with dichotic binaural presentation (DSI). A session including these two auditory tests could typically be completed in less than 45 minutes. Selected auditory processing data are available in the registered repository. Additional information on the use of the auditory processing test in PREVENT-AD and related results are detailed in two publications [34, 35].

#### 4.4 Neuroimaging

All participants were scanned on a Siemens TIM Trio 3 Tesla Magnetic Resonance Imaging (MRI) scanner at the Brain Imaging Centre of the Douglas Mental Health University Institute using a Siemens standard 12 or 32-channel coil (Siemens Medical Solutions, Erlangen, Germany). The duration of MRI sessions varied between the different visits from 0.5 to 1.5 hours and included structural and functional modalities (Figure 3A). Modalities acquired included T1-weighted, T2-weighted and Fluid-attenuated inversion recovery (FLAIR) images, diffusion MRI, arterial spin labeling (ASL), resting-state functional MRI and task functional MRI to assess episodic memory (see Table 2 for parameters of each sequence). After June 2016, new enrollees (n=48) underwent a slightly different protocol where the task fMRI acquisitions were removed and the following acquisitions added: a MP2RAGE for T1 maps, a multi-echo gradient echo for T2* maps, and a high in-plane resolution T2-weighted image to assess hippocampal subfields and brain microstructure. The 12-channel coil was replaced by a 32-channel coil for all acquisitions with this new session protocol (Figure 3B). The same images are shared in the open and the registered repositories, but images of participants presenting potentially identifying incidental findings are provided through the registered LORIS instance only.

**FIGURE 3:**
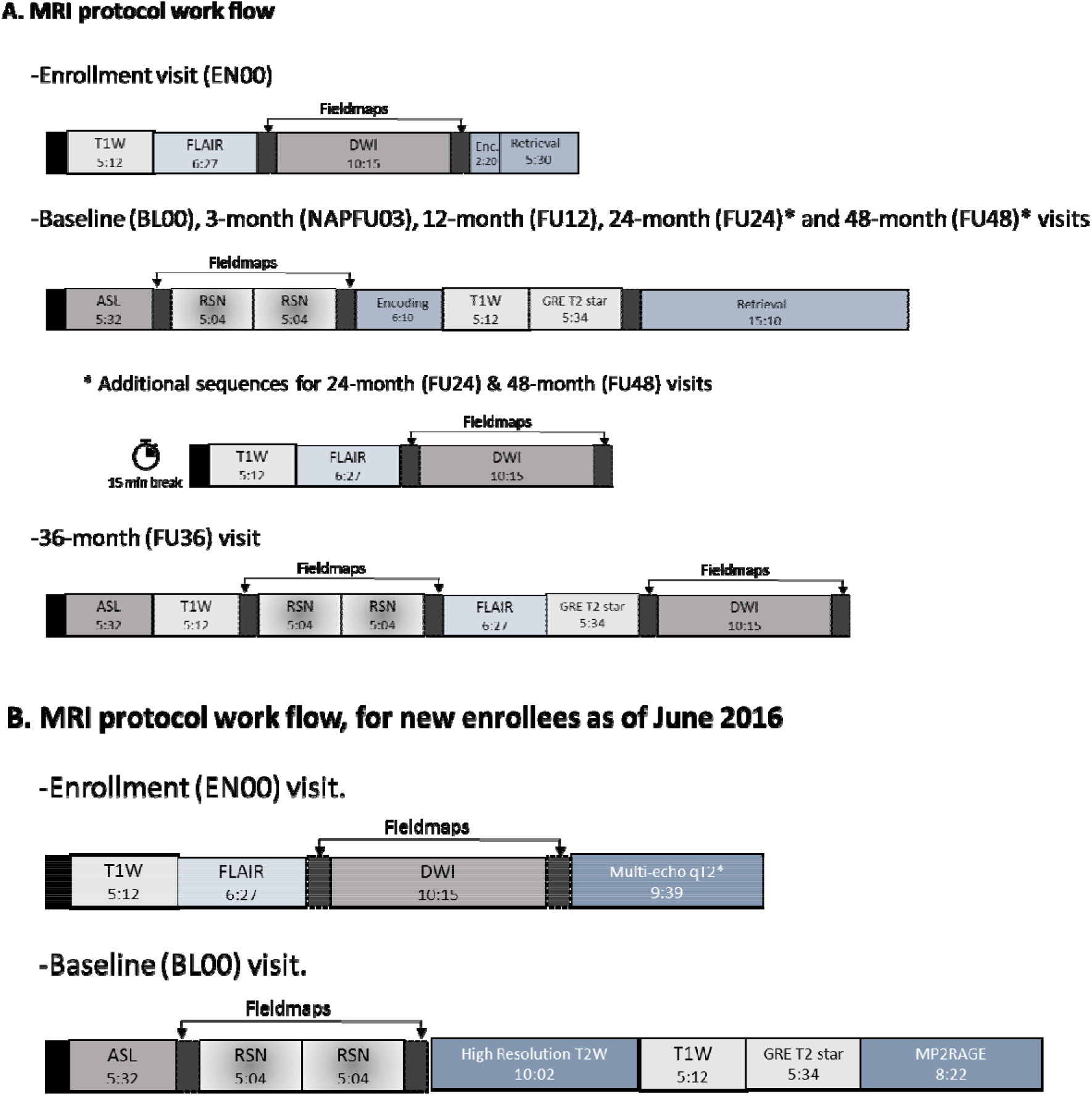
Workflow of the MRI acquisition protocol. Images from 308 scanning sessions are available in the open LORIS instance (https://openpreventad.loris.ca), while additional images of participants with incidental findings (from n=37 participants) are shared in the registered LORIS instance (https://registeredpreventad.loris.ca). **A:** The observational cohort participants (PRE) and the INTREPAD trial participants (NAP) enrolled between 2011 and May 2016 underwent the same protocol with the exception that INTREPAD trial participants performed an additional 3-month time point. The task fMRI (referred as Encoding (Enc.) and Retrieval) was performed at enrollment for practice, with the actual task performed at baseline and follow-up visits at 12, 24 and 48 months. MRI coil:12-channel. **B:** Workflow of the MRI data acquisition protocol for the observational cohort enrolled in and after June 2016 (n= 48 participants). The task fMRI protocol was replaced by a Multi-echo qT2* at enrollment and by a high-resolution T2W, GRE T2 star and a MP2RAGE at baseline. The MRI coil was upgraded to a 32-channel for this protocol. T1W = MPRAGE (Magnetization Prepared Rapid Acquisition Gradient Echo); FLAIR = FLuid Attenuated Inversion Recovery; DWI = Diffusion Weighted Imaging; ASL = Pseudo-Continuous Arterial Spin Labeling: RSN = Resting State BOLD (Blood Oxygen Level Determination); GRE T2 star = GRadient Echo T2*; Multi-echo qT2* = 12-Echo T2*; T2W = T2 -weighted.

**Table 2:** PREVENT-AD MRI parameters.

| Scan type | Sequence | Acquisition parameters | Resolution (mm3) | Scan time (min) |
| --- | --- | --- | --- | --- |
| T1-weighted anatomical | MPRAGE | 3D sagittal; TR = 2300ms; TE = 2.98ms; TI = 900ms; a = 9°; FOV = 256x240x176 mm; phase encode A-P; BW = 240Hz/px; GRAPPA 2. | 1x1x1 | 5.12 |
| Fluid attenuated T2-weighted image | FLAIR | 3D sagittal; TR = 5000ms; TE = 388ms; TI = 1800; FOV = 256x256x176 mm; phase encode A-P; BW = 781Hz/px; GRAPPA 2. | 1x1x1 | 6.27 |
| T2*-weighted anatomical | GRE | 3D transversal; TR = 650ms; TE = 20ms; a = 20°; FOV = 350x263x350 mm; phase encode R-L; BW = 200Hz/px. | 0.8x0.8x2 | 5.34 |
| Multi-echo T2*-weighted anatomical | Multi-echo GRE | 3D transversal; TR = 44ms; TE = [2.84, 6.2, 9.56, 12.92, 16.28, 19.64, 23, 26.36, 29.72, 33.08, 36.44, 39.8]ms; a = 15°; FOV = 350x263x350 mm; phase encode R-L; BW = 500Hz/px. | 1x1x1 | 9.44 |
| High-resolution T2-weighted anatomical | T2-weighted SPACE | 3D coronal; TR = 2500ms; TE = 198ms; FOV = 350x263x350 mm; phase encode R-L; GRAPPA = 2; BW = 710Hz/px. | 0.6x0.6x0.6 | 10.02 |
| T1 map | MP2RAGE | 3D sagittal; TR = 5000ms; TE = 2.91ms; TI = [700, 2500]ms; a = [4°, 5°]; FOV = 256x240x176 mm; phase encode A-P; BW = 240Hz/px; GRAPPA 2. | 1x1x1x | 8.22 |
| Diffusion-weighted imaging (DWI) | EPI | 2D transversal; TR = 9300ms; TE = 92ms; FOV = 192x192x130 mm; phase encode A-P; BW = 1628Hz/px.<br>b = [0, 1000] s/mm2 with [1, 64] directions | 2x2x2 | 10.15 |
| Perfusion imaging | Pseudo continuous-ASL (PCASL) EPI | TR = 4000ms; TR = 10ms; a = 90°; FOV = 256x256mm; 16 slices; phase encode A-P; BW = 3004Hz/px; GRAPPA 2; phase PF 7/8.<br>Label offset = 100mm; post label delay = 900ms. | 4x4x7 | 5.32 |
| resting-state functional MRI (fMRI) | EPI | 2D axial; TR = 2000ms; TE = 30 ms; a = 90°; FOV = 256x256 mm; 32 slices; phase encode A-P; BW = 2442/px. | 4x4x4 | 5.04 |
| Task functional MRI (fMRI) (Encoding and Retrieval) | EPI | 2D axial; TR = 2000ms; TE = 30 ms; a = 90°; FOV = 256x256 mm; 32 slices; phase encode A-P; BW = 2442/px. | 4x4x4 | 6.10; 15.10 |

**Legend:** TR = repetition time; TE = echo time; TI = inversion time; FOV = field of view; MPRAGE = magnetization prepared gradient echo; FLAIR = fluid attenuated inversion recovery; PCASL = pseudo-continuous arterial spin labeling

##### 4.4.1 Episodic memory task fMRI

Episodic memory tasks for object-location associations were performed by participants longitudinally, but not part of the protocol at the 36M visit. As previously mentioned, those enrolled after June 2016 did not perform this task while the existing cohort continued to complete it annually. The study design is presented in a recent publication [36]. In brief, participants were scanned as they encoded an object and its left/right spatial location on the screen. Forty-eight encoding stimuli were presented one at a time for 2000 msec with a variable inter-trial interval (ITI). A twenty-minute break followed encoding, during which time structural MRIs were acquired. After this break, participants were presented with the associative retrieval task in which they were presented with 96 objects (48 “old”-previously encoded objects; 48 “new” objects) and were asked to make a forced-choice between four-alternative answers: i) *“The object is FAMILIAR but you don’t remember the location”*; ii) *“You remember the object and it was previously on the LEFT”*; iii) *“You remember the object and it was previously on the RIGHT”*; and iv) *“The object is NEW”*. Different stimuli were employed at each visit allowing longitudinal data collection. Images used for the task were taken from a bank of standardized stimuli [37, 38]. The E-Prime program (version 2) was used to run the experimental protocol and collect behavioral data (Psychology Software Tools Inc., Pittsburgh, PA, USA).

#### 4.5 Cerebrospinal Fluid collection

Participants who consented to this procedure donated CSF samples via lumbar puncture (LP) on a separate day from the main annual visit. We link the CSF data to a specific time point as the LP procedure was performed within 6 months of this annual visit (on average 28 days after the annual visit). Given that serial LPs were initially only performed on participants enrolled in INTREPAD, the majority of the CSF samples come from INTREPAD participants (n=99 INTREPAD participants out of 160 CSF donors who consented to share data with the research community). In 2016, considering the overall success of the LP program (acceptance, tolerability, and retention through serial repetitions) serial LPs were also performed in the broader observational cohort. Therefore, some participants were enrolled in the LP protocol after their baseline visit and may have CSF data only at later time point(s). In 2017, consent for such LPs became an inclusion criterion for new participants.

LPs were performed by a neurologist (Dr. P. Rosa-Neto) with an internationally accepted procedure that typically lasted less than 15 minutes. A large-bore introducer was inserted at the L3-L4 or L4-L5 intervertebral space, after which the atraumatic Sprotte 24 ga. spinal needle was used to puncture the dura. Up to 30 ml of CSF were withdrawn in 5.0 ml polypropylene syringes. These samples were centrifuged at room temperature for 10 minutes at ~2000g, and then aliquoted in 0.5 ml polypropylene cryotubes, and quick-frozen at −80°C for long-term storage. A video describing the procedure at the StoP-AD Centre is available at https://www.youtube.com/watch?v=9kckrlBIR2E.

##### 4.5.1 CSF analysis

Biomarkers for amyloid-beta (Aβ), tau and neurodegeneration were analyzed in CSF samples. Typically, levels of Aβ_1-42_ (n=475), total tau (t-*Tau,* n=476) and phosphorylated tau (_181_p-*Tau,* n=476) were determined by enzyme-linked immunosorbent assay using Innotest technology (Fujirebio) following the European BIOMARK-APD validated and standardized protocol [39]. Additional proteins and cytokines were also analyzed as part of other subprojects and data are shared in our registered repository (ApoE ug/mL n=340, PCSK9 ng/mL n=92, G-CSF pg/mL n=321, IL-15 pg/mL n=321, IL-8 pg/mL n=321, VEGF pg/mL n=300).

### 5. Data Management

The LORIS system was customized for the PREVENT-AD program to facilitate data entry, storage, and data dissemination. Forms included customized algorithms developed for aggregating various pieces of data in a user-friendly manner. Numerous LORIS modules were also used to facilitate the curation process, including a module to track the status of the participants, a specific module on family history of AD and another one on drug compliance, for example. The document repository and data release modules facilitated the management of data distribution, documentation and access.

#### DATA RECORD

Basic demographics and longitudinal neuroimaging raw data can be found in the open LORIS repository (https://openpreventad.loris.ca), while datasets with more sensitive material such as cognitive, medical and neurosensory information, genotypes, CSF measurements, etc., are accessible to qualified researchers only at https://registeredpreventad.loris.ca. PREVENT-AD repositories are discoverable via the Canadian Open Neuroscience Platform (CONP) at https://portal.conp.ca.

Table 3 presents the list of Stage 1 PREVENT-AD shared data, their level of access, the number of participants who provided data and at which time points. MRI acquisitions available at each time point are presented in Table 4. In the registered LORIS repository, (https://registeredpreventad.loris.ca), all PREVENT-AD Stage 1 data are regrouped in 13 different CSV files accompanied by 3 text files and a detailed data dictionary. The content of each file is described in Table 5.

**Table 3:**
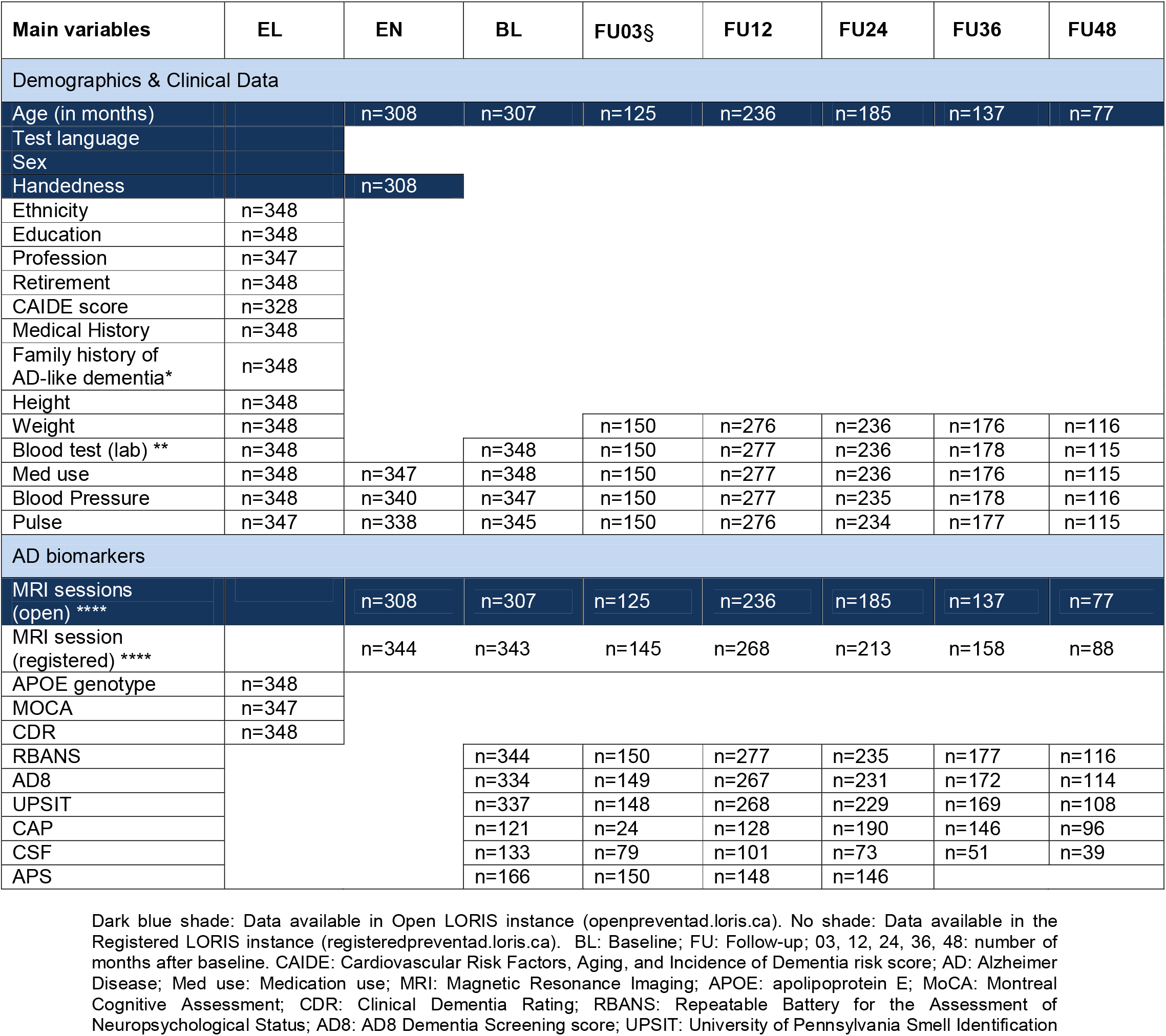
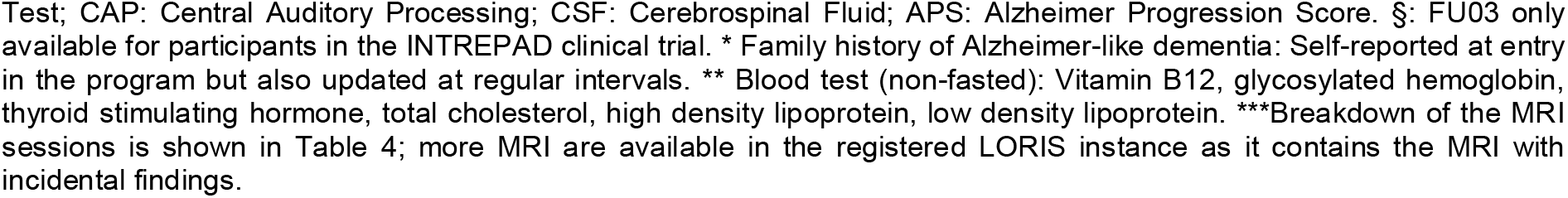
Summary of the main variables available in Stage 1 PREVENT-AD repositories at each time point (n=349).

| Main variables | EL | EN | BL | FU03§ | FU12 | FU24 | FU36 | FU48 |
| --- | --- | --- | --- | --- | --- | --- | --- | --- |
| Demographics & Clinical Data |  |  |  |  |  |  |  |  |
| Age (in months) |  | n=308 | n=307 | n=125 | n=236 | n=185 | n=137 | n=77 |
| Test language |  |  |  |  |  |  |  |  |
| Sex |  |  |  |  |  |  |  |  |
| Handedness |  | n=308 |  |  |  |  |  |  |
| Ethnicity | n=348 |  |  |  |  |  |  |  |
| Education | n=348 |  |  |  |  |  |  |  |
| Profession | n=347 |  |  |  |  |  |  |  |
| Retirement | n=348 |  |  |  |  |  |  |  |
| CAIDE score | n=328 |  |  |  |  |  |  |  |
| Medical History | n=348 |  |  |  |  |  |  |  |
| Family history of AD-like dementia* | n=348 |  |  |  |  |  |  |  |
| Height | n=348 |  |  |  |  |  |  |  |
| Weight | n=348 |  |  | n=150 | n=276 | n=236 | n=176 | n=116 |
| Blood test (lab) ** | n=348 |  | n=348 | n=150 | n=277 | n=236 | n=178 | n=115 |
| Med use | n=348 | n=347 | n=348 | n=150 | n=277 | n=236 | n=176 | n=115 |
| Blood Pressure | n=348 | n=340 | n=347 | n=150 | n=277 | n=235 | n=178 | n=116 |
| Pulse | n=347 | n=338 | n=345 | n=150 | n=276 | n=234 | n=177 | n=115 |
| AD biomarkers |  |  |  |  |  |  |  |  |
| MRI sessions (open) **** |  | n=308 | n=307 | n=125 | n=236 | n=185 | n=137 | n=77 |
| MRI session (registered) **** |  | n=344 | n=343 | n=145 | n=268 | n=213 | n=158 | n=88 |
| APOE genotype | n=348 |  |  |  |  |  |  |  |
| MOCA | n=347 |  |  |  |  |  |  |  |
| CDR | n=348 |  |  |  |  |  |  |  |
| RBANS |  |  | n=344 | n=150 | n=277 | n=235 | n=177 | n=116 |
| AD8 |  |  | n=334 | n=149 | n=267 | n=231 | n=172 | n=114 |
| UPSIT |  |  | n=337 | n=148 | n=268 | n=229 | n=169 | n=108 |
| CAP |  |  | n=121 | n=24 | n=128 | n=190 | n=146 | n=96 |
| CSF |  |  | n=133 | n=79 | n=101 | n=73 | n=51 | n=39 |
| APS |  |  | n=166 | n=150 | n=148 | n=146 |  |  |

Dark blue shade: Data available in Open LORIS instance (openpreventad.loris.ca). No shade: Data available in the Registered LORIS instance (registeredpreventad.loris.ca). BL: Baseline; FU: Follow-up; 03, 12, 24, 36, 48: number of months after baseline. CAIDE: Cardiovascular Risk Factors, Aging, and Incidence of Dementia risk score; AD: Alzheimer Disease; Med use: Medication use; MRI: Magnetic Resonance Imaging; APOE: apolipoprotein E; MoCA: Montreal Cognitive Assessment; CDR: Clinical Dementia Rating; RBANS: Repeatable Battery for the Assessment of Neuropsychological Status; AD8: AD8 Dementia Screening score; UPSIT: University of Pennsylvania Smell Identification

**Table 4.** MRI modalities available at each study visit - Stage 1.

| Scan abbreviations | EN | BL | FU03<sup>§</sup> | FU12 | FU24 | FU36 | FU48 |
| --- | --- | --- | --- | --- | --- | --- | --- |
| T1-weighted | n=308*<br>n=344 | n=307*<br>n=343 | n=125*<br>n=145 | n=236*<br>n=268 | n=185*<br>n=213 | n=137*<br>n=158 | n=77*<br>n=88 |
| ASL |  | n=307*<br>n=343 | n=124*<br>n=144 | n=236*<br>n=268 | n=184*<br>n=212 | n=70*<br>n=83 | n=77*<br>n=88 |
| DWI | n=304*<br>n=340 |  |  |  | n=184*<br>n=212 | n=132*<br>n=153 | n=73*<br>n=84 |
| FLAIR | n=308*<br>n=344 |  |  |  | n=184*<br>n=212 | n=137*<br>n=158 | n=76*<br>n=87 |
| rs-fMRI |  | n=307*<br>n=343 | n=124*<br>n=144 | n=236*<br>n=268 | n=184*<br>n=212 | n=136*<br>n=157 | n=77*<br>n=88 |
| task-encoding-BOLD | practice | n=258*<br>n=292 | n=124*<br>n=144 | n=229*<br>n=261 | n=183*<br>n=210 |  | n=76*<br>n=86 |
| task-retrieval-BOLD | practice | n=257*<br>n=291 | n=124*<br>n=144 | n=229*<br>n=261 | n=183*<br>n=210 |  | n=76*<br>n=86 |
| T2*-weighted † |  | n=307*<br>n=343 | n=125*<br>n=145 | n=236*<br>n=268 | n=184*<br>n=212 | n=136*<br>n=157 | n=77*<br>n=88 |
| MP2RAGE † |  | n=42*<br>n=44 |  | n=1*<br>n=1 |  |  |  |
| multi-echo GRE † | n=46*<br>n=48 |  |  | n=1*<br>n=1 |  |  |  |

T1-weighted = MPRAGE (Magnetization Prepared Rapid Acquisition Gradient Echo); ASL = Pseudo-Continuous Arterial Spin Labeling; DWI = Diffusion Weighted Imaging with 65 directions; FLAIR = FLuid Attenuated Inversion Recovery; rs-fMRI = Resting State functional Magnetic Resonance Imaging by Blood Oxygen Level Determination. MRIs are available in Open and Registered LORIS instance (The Registered instance also contains MRI sessions of participants presenting incidental findings that are not present in the Open instance).
\*open instance (without the scans presenting incidental findings)
§FU03 only available for people who were in the clinical trial (n=150)
† only performed on a small subset of participants (n=48)

**Table 5.** **General overview** of the content of each file, by alphabetical order, in the Stage 1 registered repository (registeredpreventad.loris.ca).

| File names | File type | Description* |
| --- | --- | --- |
| AD8_Registered_PREVENTAD | CSV | A brief informant interview (8 questions) to detect dementia [28]. |
| APS_Registered_PREVENTAD | CSV | **Alzheimer Progression Score:** composite score calculated in INTREPAD participants only [19]. |
| Auditory_processing_Registered_PREVENTAD | CSV | Central Auditory Processing, Test results from DSI and SSI-ICM [32, 33]. |
| BP_Pulse_Weight_Registered_PREVENTAD | CSV | Blood Pressure, pulse, and weight. |
| CSF_proteins_Registered_PREVENTAD | CSV | Concentration of proteins related to AD in cerebrospinal fluid (CSF). (tau, p-tau, amyloid-beta 1-42, ApoE, G-CSF, IL-15, IL-8, VEGF, PCSK9). |
| Data_Dictionary_Registered_PREVENTAD | CSV | Document providing more detail about the content of other CSV files and description of column names. |
| Demographics_Registered_PREVENTAD | CSV | General information about PREVENT-AD participants, including demographics, work related information, handedness, family history of AD and MCI converters. |
| EL_CAIDE_Registered_PREVENTAD | CSV | Cardiovascular Risk Factors, Aging, and Incidence of Dementia risk score [14]. Calculated at eligibility. |
| EL_CDR_MoCA_Registered_PREVENTAD | CSV | Cognitive Screening tests: Montreal Cognitive Assessment & Clinical Dementia Rating [15, 16]. |
| EL_Medical_history_Registered_PREVENTAD | CSV | Medical History information. |
| Genetics_Registered_PREVENTAD | CSV | Genotypes related to AD. APOE, BChE, BDNF, HMGCR, TLR4, PPP2R1A, CDK5RAP2. |
| Lab_Registered_PREVENTAD | CSV | Blood test results. Glycosylated hemoglobin, thyroid stimulating hormone, vitamin B12, total cholesterol, high density lipoprotein, low density lipoprotein. |
| List_of_participants_with_only_1_sibling | TXT | List of participants with only 1 sibling affected by Alzheimer-like dementia. |
| List_of_participants_switched_back_to_cohort | TXT | List of participants who were initially enrolled in INTREPAD trial (NAP) but failed to complete the 3-month run-in period and switched back to the cohort (PRE). |
| Med_categories | TXT | List of medications by categories. If a participant is taking a medication not in a category, it is classified as 'other'. |
| Med_use_Registered_PREVENTAD | CSV | Self-reported medication intake information |
| RBANS_Registered_PREVENTAD | CSV | Cognitive tracking test. Repeatable Battery for the Assessment of Neuropsychological Status [27]. |
| Smell_Identification_Registered_PREVENTAD | CSV | UPSIT: University of Pennsylvania smell identification test. |

*\*More details are available in the Data\_Dictionary\_Registered.csv file.*

##### Versions

Three releases were part of this Stage 1 data sharing. In April 2019, we first released data from 232 participants in the open repository (OPEN version 1.0). In August 2020, we added 76 subjects into the same open repository (OPEN version 2.0) for a total of 308 participants. We are now releasing the registered data related to 348 participants (Registered version 1.0).

#### Neuroimaging

All MRI acquisitions are available in MINC and NIfTI file formats, the latter being organized according to the Brain Imaging Data Structure (BIDS) [40]. Brain MRIs can directly be downloaded following instructions provided in both LORIS instances. MRIs are available for 308 participants in the open instance, while an additional 37 candidates with MRI presenting potentially identifying incidental findings are provided in the registered instance, for a total of 344 participants (note: 4 participants did not undergo the MRI protocol and one participant refused to share additional data in the registered repository). In the shared repositories, identifying fields (such as PREVENT-AD participant’s ID, date of birth, date of MRI, etc) were scrubbed from the MRI image headers using the DICOM Anonymization Tool (DICAT; https://github.com/aces/DICAT), while anatomical images were ‘defaced’ using the defacing algorithm developed by Fonov and Collins (2018), which has been shown to not significantly affect data processing outcomes [12]. While the code for this algorithm was slightly modified for integration into the LORIS platform, the algorithm remained unchanged. The version of the script used to deface the PREVENT-AD datasets is available in Github (https://github.com/cmadjar/Loris-MRI/blob/open_preventad_v20.1.0/uploadNeuroDB/bin/deface_minipipe.pl).

##### Task fMRI

Data were saved in .edat2 format (readable by the program only), and as text files to facilitate future data sharing. De-identification of the text files (scrubbing for dates and PREVENT-AD study IDs) was performed using a script available on Github (https://github.com/cmadjar/Loris-MRI/blob/open_preventad_v20.1.0/tools/scrub_and_relabel_task_events.pl.). De-identified data are available in both repositories.

#### TECHNICAL VALIDATION

Data were entered in LORIS in duplicate to allow detection of discrepancies between two entries of the same information and systematic corrections of mistakes by the data entry personnel. In case of significant discrepancy, source documentation was reviewed and discussed among the data entry crew and the clinical team. If needed, information was reviewed with the participants by phone, or at next follow-up. LORIS has several internal checks in place to detect any abnormalities and avoid missing data in required fields, out of range values, etc. Additional QC checks were implemented during data preparation for sharing.

##### Neuroimaging

Visual quality control of the raw anatomical images was performed by a single rater via the PREVENT-AD LORIS study interface. Quality control status, predefined comments and text comments were saved directly in LORIS. After the de-identification process, every image was visually reviewed to ensure proper defacing and the absence of any potentially identifiable information.

#### USAGE NOTE

##### Terms of use

When accessing the shared data repositories, researchers agree to a standard set of good data use practices, such as meeting ethics requirements and keeping the data secure. PREVENT-AD data must be used for neuroscience research as stipulated in the consent forms and in the Terms of Use. Authors publishing manuscripts using the PREVENT-AD Stage 1 data must name PREVENT-AD as the source of data in the abstract and or method section and cite this manuscript. The terms also include agreements on commercialization and privacy (https://openpreventad.loris.ca/login/request-account/).

For reuse of the PREVENT-AD data, researchers need to carefully read and understand the context of the data collection described in this paper and in the documentation available in the data repositories.

#### Labeling convention

The label convention used in the PREVENT-AD dataset is available when accessing the data repositories via the data dictionary.

##### Additional convention for the INTREPAD trial

Data collected for individuals enrolled in the trial are identified as such in the repositories by the prefix ‘NAP’ (e.g.: NAPFU12) in opposition to ‘PRE’ identifying the PREVENT-AD observational cohort (e.g.: PREBL00). From the shared sub-group of participants (n=349), 11 initially enrolled in INTREPAD failed to complete the first 3-months on study drug (re.: adverse events, low compliance, etc). These cases had their firsts visit labelled as ‘NAP’ and the rest as ‘PRE’ as they were switched back to the observational cohort. Any participants who stayed on the study drug for the minimum ‘run-in period’ of 3-months, kept their prefix ‘NAP’ even if the study drug was discontinued at any time between FU03 and FU24. Participants’ visits also continued to be identified as ‘NAP’ even after the end of the trial (Stage1: NAPFU36, NAPFU48 and Stage 2: up to NAPFU84).

The treatment allocation regimen (naproxen vs placebo) information is shared in the registered repository, but we suggest that data collected in the observational cohort and the trial (treated group and placebo group) can be merged for longitudinal analysis as no treatment effect was demonstrated in the trial and visit protocols were identical for all [18].

### 5. Ongoing and Future efforts

The StoP-AD Centre continues to collect data. The Stage 2 data collection regimen includes additional neuroimaging techniques, such as positron emission tomography (PET), magnetoencephalography (MEG) and a modified MRI protocol, additional lifestyle, personality traits and behavioral information as well as information on individuals who developed MCI (Pichet Binette et al., companion paper in preparation). These new acquisitions enhance the information value of the PREVENT-AD data resource and expand the number of longitudinal observations up to 84-months of follow-up. PREVENT-AD Stage 2 datasets are also being prepared to be shared with the research community. To facilitate data usage, Stage 1 and Stage 2 PREVENT-AD datasets will be shared in the same data repositories.

At the StoP-AD Centre, our goal is to continue to keep our cohort of participants engaged in our research program, carefully monitor their cognition, gather new AD biomarkers using state-of-the-art technologies and continue our involvements in the McGill University Open Science Initiatives to make data available to the greater neuroscience research community.

## ACKNOWLEDGMENTS

PREVENT-AD was launched in 2011 as a $13.5 million, 7-year public-private partnership using funds provided by McGill University, the Fonds de Recherche du Québec – Santé (FRQ-S), an unrestricted research grant from Pfizer Canada, the J.L. Levesque Foundation, the Lemaire Foundation, the Douglas Hospital Research Centre and Foundation, the Government of Canada, and the Canada Fund for Innovation. Private sector contributions are facilitated by the Development Office of the McGill University Faculty of Medicine and by the Douglas Hospital Research Centre Foundation (http://www.douglas.qc.ca/). CONP was launched in 2018 and is funded, in part, by Brain Canada. Thanks to all the financial resources.

PREVENT-AD is the result of efforts of many other co-investigators from a range of academic institutions and private corporations, as well as an extraordinarily dedicated and talented clinical and technical assistant staff, students, and post-doctoral fellows. Here is listed the entire PREVENT-AD Research Group: https://preventad.loris.ca/acknowledgements/acknowledgements.php?date=[2017-12-01]. The authors thank David Fontaine, PhD, who oversaw cognitive testing at enrollment, scored the RBANS cognitive evaluations, and administered additional cognitive evaluations when indicated. In Dr. Judes Poirier’s laboratory we wish to mention the good work that has been carried out by Anne Labonté, Doris Dea, Louise Théroux, and Cynthia Picard for CSF biomarker analyses, genotyping and more. Melissa Appleby, Laura Mahar, Miranda Tuwaig, Marie-Elyse Lafaille-Magnan, Christina Kazazian, Tanya Lee, Galina Pogossova, Renuka Giles, and Karen Wan collected cognitive and neurosensory data, assisted with the MRI sessions, entered data in LORIS and worked hard to obtain a high-quality dataset, thanks to them. Drs. Tharick Ali Pascoal, Marina Tedeshi Dauar, and Laksanun Cheewakriengkrai, dedicated time and energy to assist during LPs and performed neurological assessments with the research participants. Special thanks go to Marianne Dufour, administrative assistant, and to the clinical team; Ginette Mayrand, Joanne Frenette, Valérie Gervais, Isabelle Vallée, Rana El-Khoury, Leslie-Ann Daoust and Fabiola Ferdinand, all nurses who met with participants and were devoted to our participant’s well-being. Not to forget, everyone working on data analysis, including Jeannie-Marie Leoutsakos for the development of the APS and Melissa Savard, the PREVENT-AD data analyst. We thank Benoit Jutras, PhD, from *Université de Montreal*, for gifting us equipment to test central auditory processing. The authors would like to acknowledge the continued support and participation of the Canadian Open Neuroscience Platform (<u>CONP</u>) in making the PREVENT-AD database accessible to the scientific community. Also, the Healthy Brains for Healthy Lives (<u>HBHL</u>) initiative at McGill provided platform support for the PREVENT-AD project, through its NeuroHub IT infrastructure. We give an additional thanks to the LORIS team, at the Montreal Neurological Institute, who recently accelerated our involvement in broader data sharing, for the benefit of the community. The authors acknowledge the generosity and commitment of all research participants who volunteered for this work and placed their trusts and hope in this research program.

## AUTHORS CONTRIBUTIONS

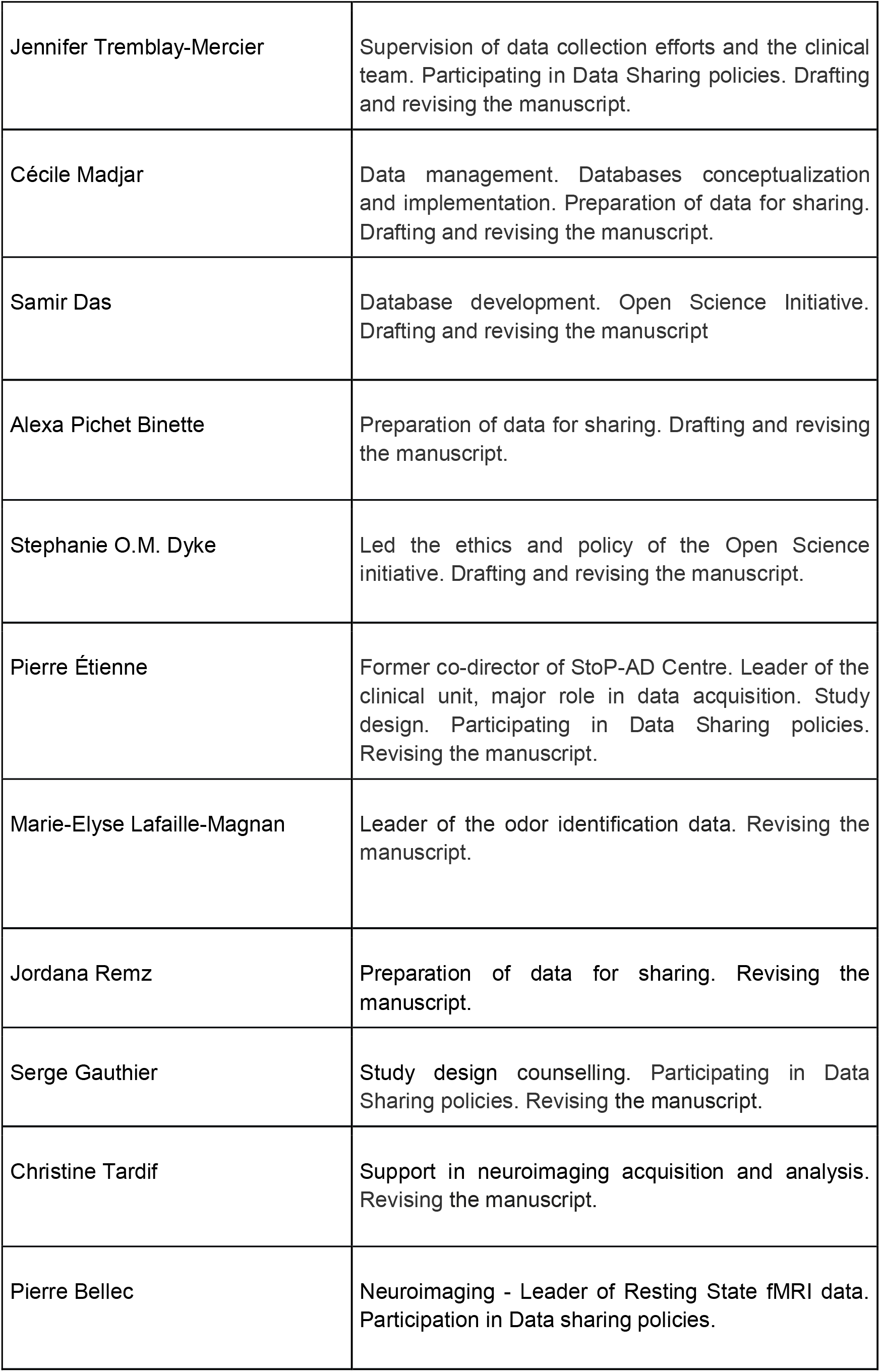
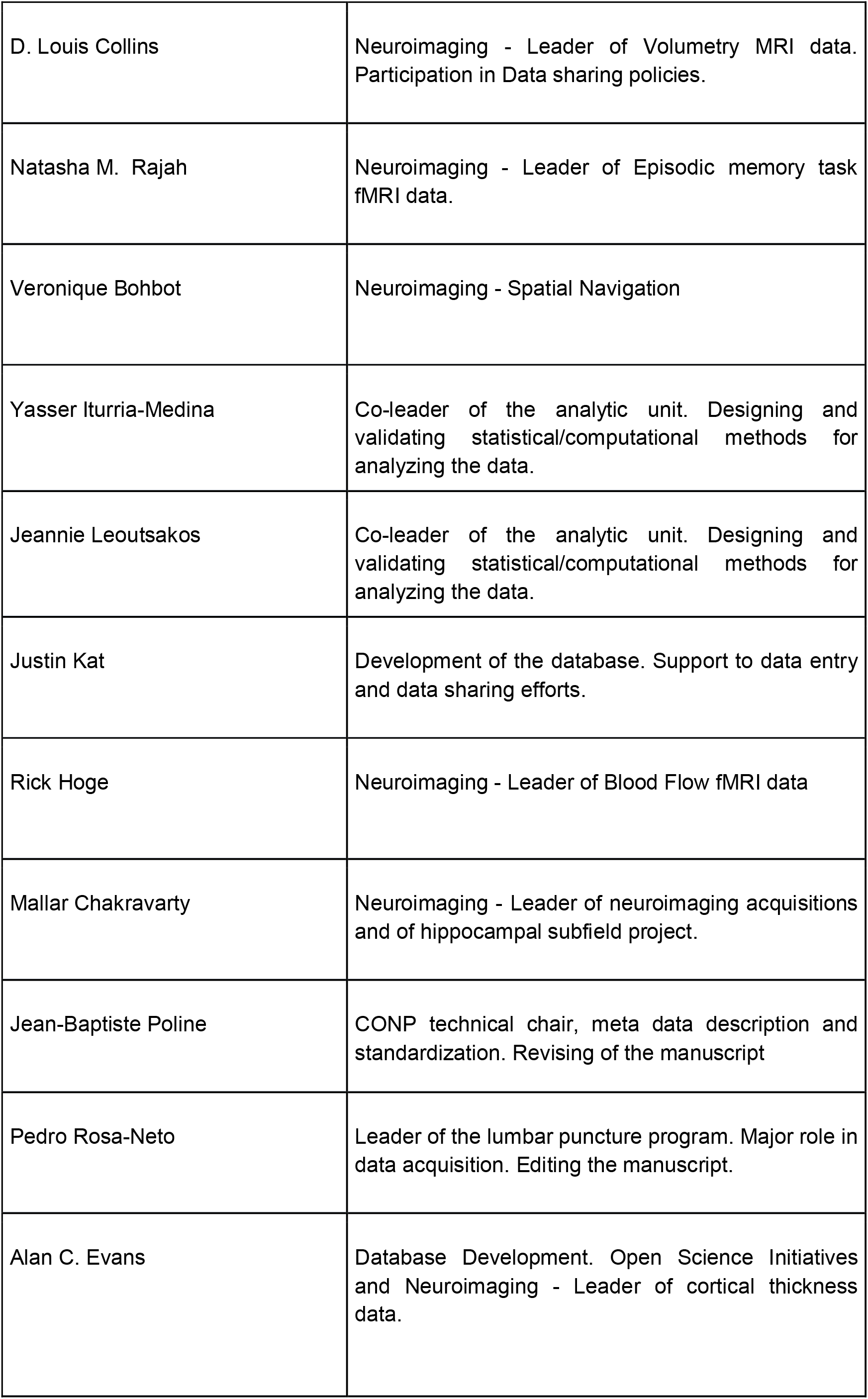
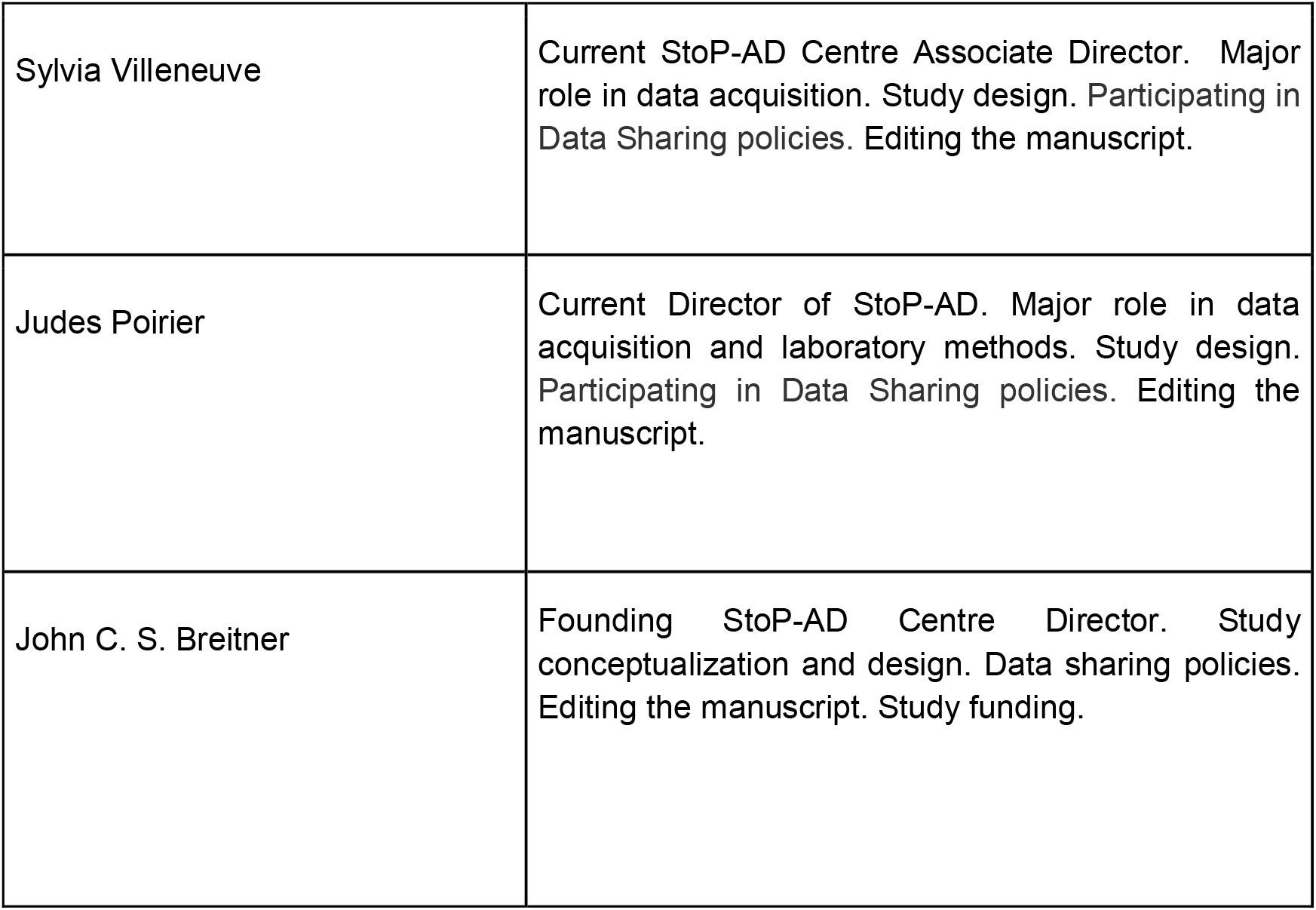

## COMPETING INTEREST

No competing interest was disclosed.

## Notes

https://openpreventad.loris.ca/

### Competing Interest Statement

The authors have declared no competing interest.

